# Long-range interhemispheric projection neurons show biased response properties and fine-scale local subnetworks in mouse visual cortex

**DOI:** 10.1101/795922

**Authors:** Kenta M. Hagihara, Ayako W. Ishikawa, Yumiko Yoshimura, Yoshiaki Tagawa, Kenichi Ohki

**Affiliations:** Department of Molecular Physiology, Kyushu University Graduate School of Medical Sciences, 3-1-1, Maidashi, Higashi-Ku, Fukuoka 812-8582, Japan; Division of Visual Information Processing, National Institute for Physiological Sciences, National Institutes of Natural Sciences, Okazaki 444-8585, Japan; Department of Physiological Sciences, The Graduate University for Advanced Studies, Okazaki 444-8585, Japan; Department of Biophysics, Kyoto University Graduate School of Science, Kitashirakawa-Oiwake-cho, Sakyo-ku, Kyoto 606-8502, Japan; Department of Physiology, Graduate School of Medical and Dental Sciences, Kagoshima University, 8-35-1, Sakuragaoka, Kagoshima 890-8544, Japan; CREST, Japan Science and Technology Agency, Kawaguchi, Saitama 332-0012, Japan; Department of Physiology, The University of Tokyo School of Medicine, 7-3-1 Hongo, Bunkyo-ku, Tokyo 113-0033, Japan; International Research Center for Neurointelligence (IRCN), The University of Tokyo, 7-3-1 Hongo, Bunkyo-ku, Tokyo 113-0033, Japan

## Abstract

Integration of information processed separately in distributed brain regions is essential for brain functions. This integration is enabled by long-range projection neurons, and further, concerted interactions between long-range projections and local microcircuits are crucial. It is not well known, however, how this interaction is implemented in cortical circuits. Here, to decipher this logic, using callosal projection neurons (CPNs) as a model of long-range projections, we found that CPNs exhibited distinct response properties and fine-scale local connectivity patterns. *In vivo* 2-photon calcium imaging revealed that CPNs showed a higher ipsilateral eye (with respect to their somata) preference, and that CPN pairs showed stronger signal/noise correlation than random pairs. Slice recordings showed CPNs were preferentially connected to CPNs, demonstrating the existence of projection target-dependent fine-scale subnetworks. Collectively, our results suggest that long-range projection target predicts response properties and local connectivity of cortical projection neurons.

## Introduction

The cerebral cortex processes information arriving from the external world in a distributed manner. To achieve this, there must be biophysical substrates enabling proper routing, local computation, and integration of information. Integration of distributed information is mediated by cortical projection neurons; thus, the functional organization of these neurons have been extensively studied (Glickfeld et al., 2013; Kim et al., 2015; Lur et al., 2016; Movshon and Newsome, 1996; Sato and Svoboda, 2010; Yamashita et al., 2013). Such studies have revealed that as a general principle, neurons projecting to different areas encode different features of the sensory input. In addition, fine-scale local network motifs (Kampa et al., 2006; Yoshimura et al., 2005) implicated in selective local computation have been identified. Recent work has found that cortical neurons that do not share their projection targets are rarely connected to each other, suggesting the existence of strongly segregated local network motifs suited for highly selective information transmission (Brown and Hestrin, 2009; Kim et al., 2018). However, the organizational principles linking projection targets to functional properties and fine-scale local networks have not been fully addressed. More specifically, whether the strongly segregated local network is the general motif for all the long-range projection populations is an open question to be addressed.

Callosal projection neurons (CPNs) are essential for integrating information processed in separate hemispheres (Hubel and Wiesel, 1967; Van Essen et al., 1982), and thus serve as an ideal model to study the functional organization of long-range projection neurons. Although many lesion and inactivation studies have been performed, to what extent callosal inputs influence the visual response properties of primary visual cortex (V1) neurons is still controversial (Berlucchi and Rizzolatti, 1968; Cerri et al., 2010; Dehmel and Lowel, 2014; Minciacchi and Antonini, 1984; Payne et al., 1980; Restani et al., 2009; Schmidt et al., 2010; Wunderle et al., 2013; Zhao et al., 2013). Moreover, due to the lack of a method enabling selective *in vivo* recordings from CPNs and non-CPNs, the specific visual information encoded and transmitted by individual CPNs remains largely unknown.

In higher mammals such as cats and monkeys, the inputs from each eye are spatially segregated in V1 as ocular dominance columns (Van Essen et al., 1992; Wiesel and Hubel, 1965). On the other hand, rodents, especially mice have no ocular dominance columns ((Antonini et al., 1999; Drager, 1974; Gordon and Stryker, 1996; Mrsic-Flogel et al., 2007; Scholl et al., 2015) but see (Laing et al., 2015) for results in the Long Evans rat;) rather, functionally distinct neuronal populations are spatially organized in a “salt and pepper” manner (Ohki and Reid, 2007). Thus, to assess the relationships between CPN response properties and local connection patterns, it is crucial to assess neuronal responses and local connectivity at single-cell resolution and to directly compare the functional organization of CPNs to surrounding non-CPNs. Here, we achieve this by combining retrograde-labeling of CPNs with *in vivo* two-photon calcium imaging and *ex vivo* slice recordings.

## Results

To compare the visual response properties of CPNs and non-CPNs in layer 2/3 (L2/3) of mouse visual cortex, we performed *in vivo* two-photon calcium imaging (Ohki et al., 2005). Using visual stimulation to each eye, we first assessed the ocular dominance response properties of CPNs and non-CPNs. The fluorescent retrograde tracer cholera toxin B subunit conjugated to the fluorophore Alexa 555 (CTB555) was used to label CPNs (**Figure 1A, B, Methods)**. As Oregon Green BAPTA-1 (OGB1) injection was also performed to largely cover the CTB555-positive region along the medial–lateral axis, our recordings equally contained binocular regions of V1 and lateral secondary visual cortex (more specifically LM (Wang and Burkhalter, 2007)). We imaged 1,866 CPNs and 11,399 non-CPNs from 7 mice. There was no difference in the fraction of visually responsive CPNs and non-CPNs (31.3% ± 1.8% vs. 28.9% ± 2.2%, *P* = 0.30, Wilcoxon signed-rank test, *n* = 7 mice). Based on fluorescence signal changes in response to unilateral visual inputs, we calculated the ocular dominance (OD) score (see **Methods** and (Mrsic-Flogel et al., 2007)) for all visually responsive neurons. Surprisingly, some CPNs showed exclusive ipsilateral eye preference (**Figure 1C**). While OD scores varied extensively for both CPNs and non-CPNs, the proportion of neurons exhibiting a high OD score (with ipsilateral eye preference) was larger in CPNs than non-CPNs (**Figure 1D**). Indeed, a significantly larger fraction of CPNs had OD scores >0.8 compared to non-CPNs (**Figure 1E**, *P* = 0.016, Wilcoxon signed-rank test, *n* = 7 mice), while there was no significant difference in the fraction of OD scores <0.2 (*P* = 0.47). As previously reported in mice (Antonini et al., 1999; Drager, 1974; Gordon and Stryker, 1996; Mrsic-Flogel et al., 2007; Scholl et al., 2015), we did not observe any evidence for spatial clustering or microstructure related to eye preference or to CPNs in C57BL/6 mice (**Figure S3**). Both CPNs and non-CPNs showed similar tuning properties for drifting gratings, with only slightly greater orientation selectivity in CPNs (**Figure S4A-C**). These results suggest that CPNs have somewhat distinct visual response properties from non-CPNs.

**Figure 1.**
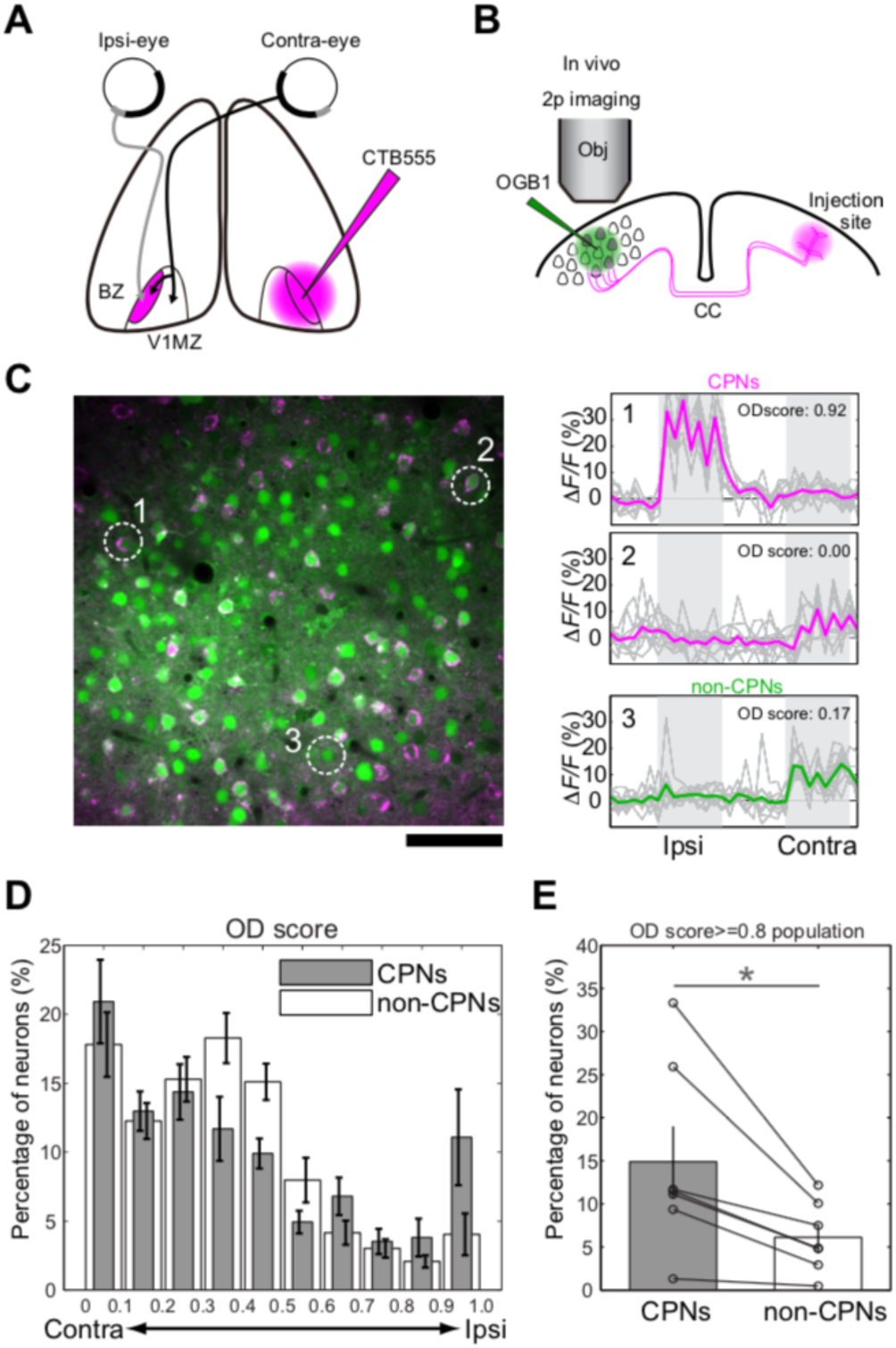
Ocular dominance response properties of CPNs and non-CPNs. **A.** Schematic drawing of CPNs labeling with retrograde tracer CTB555. BZ: binocular zone; V1MZ: primary visual cortex monocular zone. **B.** Schematic drawing of *in vivo* two-photon calcium imaging of CTB555 labeled CPNs and surrounding non-CPNs. **C.** (Left) Image of OGB1-labeled (green) and retrogradely CTB555-labeled (magenta) callosal projection neurons (CPNs) and non-CPNs in the binocular zone (BZ). Scale bar: 20 μm. (Right) Representative calcium traces from CPNs (#1 and #2) and a non-CPN (#3) shown in the left panel. **D.** Distribution of ocular dominance (OD) scores for CPNs and non-CPNs. **E.** Proportion of neurons with an OD score over 0.8. * *P* < 0.05; (n = 7 mice, Wilcoxon signed-rank test) See also **Figure S1 and S2**

Although CPNs are not organized into columns or spatial clusters in mice (**Figure S3**), it is still possible that they form functional subnetworks. To test this possibility, we analyzed the signal and noise correlations of CPN and non-CPN activity (**Figure 2, S4D, E**). Both signal and noise correlations were significantly stronger between CPN pairs than between CPN/non-CPN pairs and non-CPN/non-CPN pairs (**Figure 2B**, CPN/CPN vs. CPN/non-CPN, *P* < 0.001; CPN/CPN vs. non-CPN/non-CPN, *P <* 0.001; **Figure 2E**, CPN/CPN vs. CPN/non-CPN, *P <* 0.001; CPN/CPN vs. non-CPN/non-CPN, *P <* 0.001; *n* = 62 planes from 7 mice; Friedman test followed by Tukey-Kramer method). These differences also held true when we analyzed neuron pairs of similar soma–soma distance (**Figure 2C** and **F**). Thus, CPNs are likely to form local functional subnetworks that are organized independently of cell body location.

**Figure 2.**
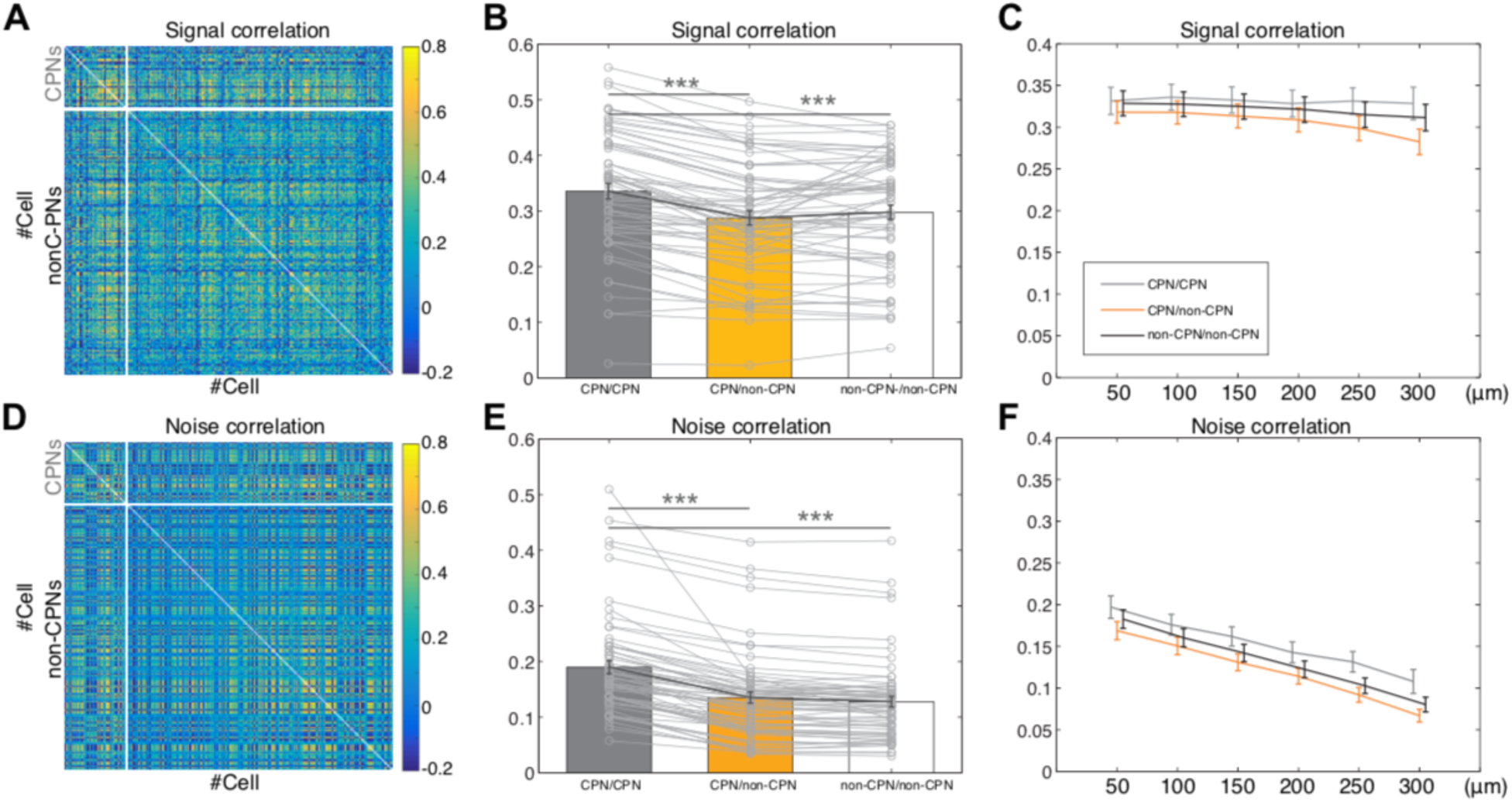
Correlation structure of CPNs and non-CPNs. **A, D**. Representative signal (**A**) and noise (**D**) correlation matrices of neurons from a single imaging plane. The order of neurons is sorted so that CPNs are located at the top and non-CPNs at the bottom. **B, E**. Means of signal (**B**) and noise (**E**) correlation values from each plane. CPN/CPN pairs show both higher signal and noise correlation coefficient values than CPN/non-CPN and non-CPN/non-CPN pairs. **C, F**. Signal (**C**) and noise (**F**) correlations of CPN/CPN (gray), CPN/non-CPN (orange), and non-CPN/non-CPN (black) pairs with respect to cortical distance. ***: *P* < 1.0 × 10^−3^; (Friedman test followed by Tukey-Kramer). See also **Figure S3 and S4**

Pairs of V1 neurons showing stronger signal/noise correlation show higher probability of synaptic connections (Cossell et al., 2015; Ko et al., 2011). To directly test whether CPN pairs show higher connection probability, we conducted dual whole-cell patch-clamp recordings from CPNs and/or non-CPNs in slices of the hemisphere contralateral to the CTB injection site at 4–8 days post-injection, specifically targeting the most densely CTB-labeled area in L2/3 (**Figure 3A, B**). The probability of monosynaptic excitatory connections between CPN/CPN pairs (30%) was significantly higher than that between CPN/non-CPN pairs (12.2%, *P* = 0.046, Fisher’s exact test, **Figure 3C**). The probability of connections between non-CPN/non-CPN pairs (19.1%) also tended to be lower than that of CPN/CPN pairs, although the difference did not reach significance. Reciprocal connections were rare between CPN/CPN pairs (1/50), CPN/non-CPN pairs (0/41), and non-CPN/non-CPN pairs (0/42). The connection probability for all recorded pairs (21.1%, 28/133) was in agreement with that for randomly sampled pairs in L2/3 visual cortex (Cossell et al., 2015; Ko et al., 2011). The amplitudes of EPSCs recorded from connected CPN/CPN pairs were comparable to those measured from connected CPN/non-CPN and connected non-CPN/non-CPN pairs (*P* = 0.95, Kruskal–Wallis test, **Figure 3D**). Also, there were no statistical differences in intrinsic membrane properties between CPNs and non-CPNs (**Figure S5)**. Overall, these results demonstrate that L2/3 neurons sharing long-range projection targets form functional subnetworks defined by local connectivity.

**Figure 3.**
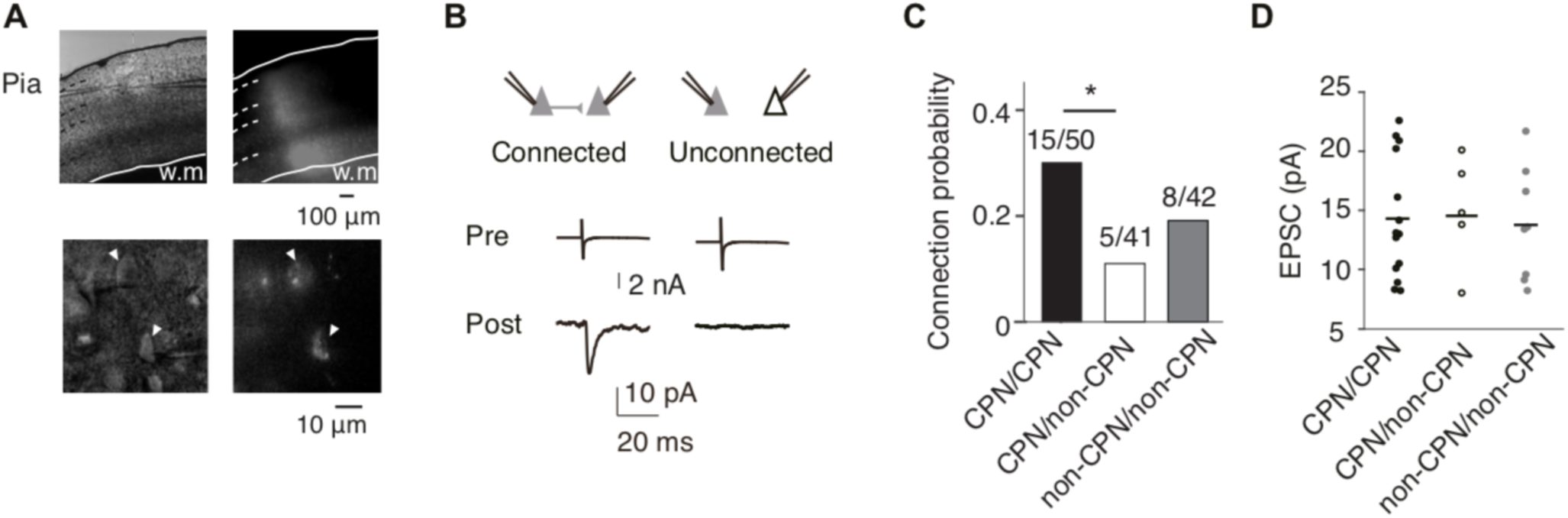
Local synaptic connectivity of CPNs. **A.** Top-left: Differential interference contrast image showing a recording electrode position in L2/3. Top-right: Corresponding image of CTB fluorescence in a coronal slice of primary visual cortex containing the binocular zone. Bottom-left: Higher magnification of the interference contrast image showing recorded neurons. Bottom-right: Corresponding high-magnification fluorescent image of CTB-labeled CPNs. White triangles mark CTB-labeled recorded neurons. **B.** Example averaged traces of presynaptic action currents evoked by depolarizing voltage pulses and resultant EPSCs in postsynaptic cells. **C.** Connection probabilities of CPN/CPN, CPN/non-CPN and non-CPN/non-CPN pairs. The connection probability of CPN/CPN pairs was significantly higher than that of CPN/non-CPN pairs (^*^*P* < 0.05 from Fisher’s exact test). The numbers of recorded and connected cell pairs are shown above each bar. **D.** Amplitudes of evoked EPSCs obtained from connected neuron pairs. Black lines indicate medians. See also **Figure S5**

## Discussion

In the current study, we used CPNs as a model to investigate the functional properties and local circuit organization of long-range projection neurons in the cortex. How CPNs and other projection neurons in the early visual cortex distribute visual feature information in a target-specific manner is an intriguing question. Our data showing stronger signal/noise correlations (**Figure 2**) and local connectivity among CPNs (**Figure 3**) suggest the existence of target-specific local functional subnetworks in L2/3. It has been reported that local subnetworks of neurons sharing similar visual properties emerge through activity-dependent synaptic plasticity (Ishikawa et al., 2014, 2018; Ko et al., 2013). Our data extend these findings by showing that long-range projection specificity is another factor determining local subnetwork organization in L2/3, a notion also supported by a recent study that assessed V1 neurons projecting to different higher visual cortices (Kim et al., 2018). In contrast to the strongly segregated subnetworks found in neurons projecting to different higher-order visual areas within the same hemisphere (Kim et al., 2018), moderately segregated subnetworks between CPNs and non-CPNs reported in the current study may allow for interactions between the subnetworks of CPNs and other neurons in the local network. Overall, this projection target-dependent segregation of local subnetworks may be a general principle for intermingled projection neurons in the mouse cortex. That said, different levels of segregation in distinct projection populations are of note. The extent of segregation would define the selectivity of information to be extracted locally and then transmitted to the projection target.

We found that a larger fraction of CPNs than non-CPNs preferentially responded to ipsilateral eye input. Some of them even showed exclusive ipsilateral eye preference (**Figure 1**). Considering that most of the retino-thalamic afferents cross in the chiasm (Lund, 1965), and that most of the lateral geniculate neurons show binocular response in rodents (Howarth et al., 2014), this strong ipsilateral selectivity found in CPNs was rather unexpected. This ipsilaterally-biased CPN population may serve to duplicate visual information conveyed by the thalamo-cortical projection, and send a copy of that information to the other hemisphere (Laing et al., 2015). Thus, callosal connections may function as a back-up system for the thalamo-cortical projection (Li et al., 2016). Different from other projection neurons in V1 (Glickfeld et al., 2013), CPNs show similar SPF/TF tuning as non-CPNs (**Figure S5**). This property may also be useful for the putative back-up function. In contrast to our findings, a recent study (Lee et al., 2019) did not find a strongly ipsilaterally biased population (to the hemisphere where CPN somata are located) among callosal inputs to L2/3 V1 neurons (contralateral to the callosal recipient neurons), raising the possibility that the ipsilaterally biased callosal population selectively targets contralateral L5 neurons, which receive the largest fraction of callosal inputs (Petreanu et al., 2007), whereas majority of callosal neurons, which show contralateral eye (to the hemisphere where CPN somata are located) preference, might target contralateral L2/3 neurons.

Several recent studies have shown that V1 neurons projecting to different higher-order visual cortices (Glickfeld et al., 2013; Matsui and Ohki, 2013) and subcortical areas show distinct response properties (Kim et al., 2015; Lur et al., 2016). However, the extent of overlap through collateral projections (Han et al., 2018) is not known. For instance, CPNs could overlap with other specific projecting population (Yamashita et al., 2013; Yamashita et al., 2018). Thus, comprehensive characterization of the relationships among projection target, response properties, molecular identity, and local connectivity is required for a better understanding of the circuit logic in the early visual cortex, which functions as a central hub to process visual information.

## Acknowledgements

We thank D. Hillier (Friedrich Miescher Institute), T. Kanamori (University of Basel), E.M.M. Meyer (FMI), and R.K. Morikawa (FMI) for reading and commenting on earlier versions of the manuscript; all of the members of Ohki laboratory for support and discussion; A. Honda (Ohki lab) for histology; Y. Sono (Ohki lab) for animal care; the Research Support Center, Graduate School of Medical Sciences, Kyushu University for technical support. This work was supported by grants from CREST-JST (to K.O. and Y.T.), JSPS KAKENHI (Grant number 25221001, 25117004, 19H01006 to K.O., 23500388, 16K06992 to Y.T., 17K14942 to A.W.I), “Neural Diversity and Neocortical Organization” (23123508 and 25123707 to Y.T.) and “Dynamic regulation of brain function by Scrap & Build system” (16H06460 to Y.Y., 17H05745 and 19H04756 to Y. T.). A part of this work was carried out under the Brain/MINDS by the MEXT of Japan. K.M.H was supported by Takeda Science Foundation. Y.T. was supported by Astellas Foundation for Research on metabolic Disorders, the Kodama Memorial Fund for Medical Research, and the Novartis Foundation (Japan) for the Promotion of Science.

## Author Contributions

K.M.H. and Y.T. initially conceived the research. K.M.H. designed and performed imaging experiments and analyzed the data. K.O. supervised imaging experiments, data analysis and interpretation of the data. A.W.I designed and performed slice experiments and analyzed the data. Y.Y. supervised slice experiments, data analysis and interpretation of the data. K.M.H., Y.T., A.W.I. wrote the manuscript. K.O. and Y.Y commented on the manuscript.

The authors declare no competing interests.

## Materials and Methods

### LEAD CONTACT AND MATERIALS AVAILABILITY

Further information and requests for resources and reagents should be directed to and will be fulfilled by the Lead Contact, Kenta M. Hagihara.

### EXPERIMENTALMODEL AND SUBJECTDETAILS

All of the experiments were performed in accordance with the institutional animal welfare guidelines of the Animal Care and Use Committee of Kyushu University, Kyoto University and National Institute for Physiological Sciences, and they were approved by the Ethical Committee of Kyushu University, the local committee for handling experimental animals in the Graduate School of Science, Kyoto University, and National Institute for Physiological Sciences.

## METHOD DETAILS

### Stereotactic retrograde tracer injection for imaging experiments and histology

Mice (C57BL/6, both males and females, aged at postnatal day 60-90) were housed in the temperature-controlled animal room with 12h/12h light/dark cycle. Cholera Toxin Subunit B Alexa Fluor 555 or 488 Conjugate (CTB555 or CTB488, 1.0%, weight/volume, Invitrogen) and Green Retrobeads™ IX (Green Beads, Lumafluor Inc.) were used as retrograde tracers (Conte et al., 2009; Li et al., 2015). The mice were subjected to stereotactic tracer injections using pulled glass micropipettes. The mice were anesthetized with 440 mg/kg chloral hydrate (Tokyo Chemical Industry) by intraperitoneal injection. The mice were then fixed to a small-animal stereotactic device (Narishige). The head skin was cut at the midline, and the periosteum was removed using a surgical knife. The skull was thinned with a drill, and a small craniotomy was made using a 30-gauge needle. The tracer was injected through a pulled glass micropipette connected to a Hamilton syringe (Hamilton Company), which was pumped using a syringe pump device (World Precision Instruments). The stereotactic injections were administered at BZ or the secondary motor cortex (M2). The stereotaxic coordinates for BZ injection were 3.5 mm lateral of the midline and 2.0 mm in front of the anterior margin of the transverse sinus. For M2 injections, 0.5 mm lateral and 0.5 mm anterior to the bregma. The tracer solution was injected at a rate of 0.05–0.1 μl/min at a volume of 2.0 μl for BZ and 0.5 μl for M2, and after the injection, the pipette was held in place for an additional 10 min before removal. After the removal of the micropipette, the skin incision was sutured. The post-injection animals were bred normally for 5–14 days before imaging or histology experiments. Fluorescent histological images were acquired using a confocal laser-scanning microscope (LSM700, Zeiss).

In the experiment described in **Figure S2**, a cocktail of Green Beads and CTB555 (1.0%) was used. Given the efficiency (0-1) of Green Beads and of CTB555 as *a* and *b*, respectively, and the fraction of CPNs as *n*, the fraction of CPNs labeled with both Green Beads and CTB555 would be *n*a*b*. Considering that neurons labeled with Green Beads were also mostly labeled with CTB555, here n*\*a*b* can be regarded as near *n*a* and thus *b* can be regarded as 1. Thus, in this system, the efficiency of CTB555 can be regarded as almost 100% and neurons not labeled with CTB555 can be regarded as non-CPNs.

### Stereotactic retrograde tracer injection for slice experiments

For retrograde labeling of CPNs, 28–32-day old C57BL/6 mice of either sex were anesthetized with propofol (0.2%, 100 ml/kg ip, Maruishi) and then fixed to a small-animal stereotactic device. CTB488 or CTB555 (300–500 nl) was pressure-injected (Toohey Spritzer microinjector, Toohey Company) into the binocular zone (BZ) of the visual cortex contralateral to the eventual recording site (3.5 mm lateral to the midline, 1.5 mm rostral to the anterior margin of the transverse sinus, 300–600 μm in depth) with pulled glass pipettes (tip diameter 10–20 μm, Narishige). The pipette was held in place for 10 min before and after injection. After removal of the pipette, the skin incision was sutured and mice were allowed to recover for 4–8 days before recording.

### Animal preparation and surgery for in vivo calcium imaging

5-14 days after CTB555 injection, the animals were prepared for *in vivo* calcium imaging as previously described (Hagihara et al., 2015; Ohki and Reid, 2014). In brief, mice were anesthetized with isoflurane, then a custom-made metal plate was mounted on the skull, and a craniotomy was carefully performed on the other hemisphere above the area where fluorescence from CTB555 was clearly observed through the skull. We dissolved 0.8 mM Oregon Green 488 BAPTA-1 AM (OGB1) in DMSO with 20% pluronic acid and mixed it with ACSF containing 0.05 mM Alexa594 (all obtained from Invitrogen). A glass pipette (3–5 μm tip diameter) was filled with this solution and inserted into the cortex, and OGB1 and Alexa were pressure-ejected from the pipette (−2 psi for 300–500 ms, 10–20 times). After confirming loading, the craniotomy was sealed with a cover glass. 281.6 μm×281.6 μm area was imaged using a two-photon microscope (Nikon A1MP, Nikon), which was equipped with a mode-locked Ti:sapphire laser (MaiTai Deep See, Spectra Physics) at 2 Hz with 512×512 pixels (0.55 μm/pixel). The excitation light was focused with a 25× Nikon (NA: 1.10) PlanApo objective. The average power delivered to the brain was < 20 mW, depending on the depth of focus. OGB1 and CTB555 were excited at 1000 nm except for the experiment using 920 nm described in **Figure S1**. The emission filters were 517–567 nm for OGB-1 and 600–650 nm for CTB555.

### Visual stimulation and image data acquisition

The visual stimulation for studying the binocular response property was performed using two LEDs controlled by Arduino (Smart Projects). To avoid contamination to the other eye, LEDs were directly attached to the eyes with black silicon. Intensity was set as 40 cd/m^2^. Each stimulus started with a blank period (4s), which was followed by visual stimulation (4s). Each stimulus was consisted of 4 rounds of alternative 0.5 seconds of ON and OFF periods. We used 410 nm LED stimuli, based on the report that all mouse photoreceptors display similar sensitivity to spectrum around 400 nm (Govardovskii et al., 2000). Our measures of sensitivity are thus not biased towards responses originating from any specific photoreceptor. In particular, the dorsal-ventral gradient in cone-opsin expression in the mouse retina (Applebury et al., 2000; Sterratt et al., 2013) is not likely to have influence with our measures of relative sensitivity. To study orientation and direction selectivity, drifting square-wave gratings (100% contrast, 2 Hz temporal frequency) were presented in 12 directions of motion with 30 degree steps on a 27-inch LCD monitor (Samsung, Hwaseong, South Korea) placed in 20 cm distance from eyes. We positioned the center of the monitor to cover the vertical meridian. The drifting gratings visual stimulus sets were generated using custom-made software written in PsychoPy (Peirce, 2007). The spatial frequency was set at 0.03 cycles per degree. Each stimulus started with a blank period of uniform gray with the same mean luminance of gratings (4s), which was followed by visual stimulation (4s). For mapping of spatial frequency (SPF) and temporal frequency (TF) tunings, drifting sine-wave gratings (100% contrast) were used. For SPF mapping experiments, sine-wave gratings with six SPFs between 0.01 cpd and 0.32 cpd in octave steps, drifting at 2 Hz were used. For TF mapping experiments, sine-wave gratings having 0.04 cpd and drifting at 5 different TFs between 0.5 and 8 Hz in octave steps were used. Each stimulus started with a blank period of uniform gray (4 s) followed by the same period of visual stimulation during vertical and horizontal gratings were presented for 1 s for each of four directions (0°, 180°, 90°, and 270° in that order). These 3 drifting grating stimulus sets were repeated 10 times.

### Slice experiments

Coronal slices (300 µm thick) containing visual cortex were prepared from mice at postnatal day 32–36 following deep anesthesia with isoflurane. At this age, binocular matching of orientation preference in V1 neurons reaches the adult level, suggesting that binocular visual functions in V1 are mature (Wang et al., 2010). Prior to recording, slices were incubated for 1 h in oxygenated (95% O_2_ and 5% CO_2_) normal artificial cerebrospinal fluid (ACSF) containing (in mM) 126 NaCl, 3 KCl, 1.3 MgSO_4_, 2.4 CaCl_2_, 1.2 NaH_2_PO_4_, 26 NaHCO_3_, and 10 glucose at 33°C as described previously (Ishikawa et al., 2014). Pyramidal neurons in L2/3 were targeted by patch pipettes under fluorescent and infrared differential interference contrast (DIC) optics with a 60X water immersion objective (BX51, Olympus, Tokyo, Japan, **Figure 3A**). Cells retrogradely labeled with CTB488 or CTB555 were identified by epifluorescence with appropriate filter sets (U-MWIB3: excitation 479–495 nm and emission 510IF nm for CTB488; U-MWIG3: excitation 530–550 nm and emission 575IF nm for CTB555). We performed whole-cell recordings from pairs of L2/3 pyramidal neurons with somata separated by <70 μm. Cell bodies of the recorded neurons were located >50 μm below the cut surface of the slice. The patch pipettes (4–6 MΩ) were filled with an internal solution containing (in mM) 130 K-gluconate, 8 KCl, 1 MgCl_2_, 0.6 EGTA, 10 HEPES, 3 MgATP, 0.5 Na_2_GTP and 10 Na-phosphocreatine (pH 7.3, adjusted using KOH). The presence of synaptic connections between recorded neuron pairs was tested by applying brief depolarizing voltage pulses (duration, 1–2 ms) to evoke an action potential in one cell (≥ 50 trials) and recording excitatory postsynaptic currents (EPSCs) from the other cell. The membrane potential of recorded cells was held at the reversal potential of IPSCs (−70 mV). Recordings were made using a Multiclamp 700B amplifier, Digidata 1440A converter, and pCamp10 software (all from Molecular Devices).

Data were digitized at 10 kHz, filtered at 1 kHz, and analyzed using custom-made programs written in Matlab. To obtain the amplitude of EPSCs, we averaged the peak values of individual EPSCs after excluding failures. The baseline current was defined as the averaged current in a 5-ms window before application of depolarizing voltage pulses to the presynaptic cell. The peak EPSC was defined as the peak current minus the baseline. We selected cells with seal resistance >1 GΩ and series resistance <35 MΩ for analysis. For measuring intrinsic membrane properties of neurons, 1-s step currents from −100 pA to 500 pA in 25 pA increments were injected in current clamp mode. The current injection experiments were conducted within 5 min after establishing the whole-cell patch-clamp configuration.

## QUANTIFICATION AND STATISTICAL ANALYSIS

### Data analysis of visual response

The images were analyzed by using our custom-written in-house software running on Matlab (Mathworks) as previously described (Hagihara et al., 2015; Ohki et al., 2005). In brief, the cell outlines were automatically identified using template matching. The identified cell outlines were visually inspected, and the rare but clear errors were manually corrected. The time courses of individual cells were extracted by summing the pixel values within the cell outlines. Slow drift of the time courses over minutes was removed by applying a low-cut filter (cut-off, 2–4 min). The baseline (F) for each trial with each cell was calculated by averaging the values of the last 0.5 s of the blank periods for all stimuli. Neuropil contamination was removed as previously described (Hagihara et al., 2015; Kerlin et al., 2010). Visually responsive cells were defined by ANOVA (*P* < 0.01) across blank and 2 eyes (for ocular dominance stimuli), across blank and 12 direction periods (for orientation/direction stimuli), across blank and 6 SPF/TF periods (for SPF/TF stimuli), where ΔF/F>2%. Instead of *P* < 0.05, *P* < 0.01 was used here, to reduce false positives due to large sample size of neurons.

For responsive cells with ocular dominance stimuli, ODscore (Mrsic-Flogel et al., 2007) was defined as follows: (response to ipsilateral-eye)/(response to ipsilateral-eye) + (response to contralateral-eye). For responsive cells with drifting grating stimuli, as an index for orientation selectivity tuning, gOSI was calculated. This value is equivalent to :((Sigma R(θ_i_)sin(2θi))^2 + (Sigma R(θi)cos(2θi))^2)^(1/2)/Sigma R(θi), where θi is the orientation of each stimulus and R(θ_i_) is the response to that stimulus. For direction selectivity, DI was calculated as :1 – (response to null direction)/(response to preferred direction). The preferred orientation and direction were defined by vector averaging of responses (ΔF/F) to all orientations and directions, respectively. For correlation analysis, Pearson’s correlation coefficient was calculated to obtain pair-wises response correlation. Signal correlation was calculated as the correlation coefficient between the trial-averaged traces in responses to the ocular dominance stimulus. Noise correlation was obtained by subtracting the trial-averaged traces from the traces to each trial, and then calculating the correlation coefficient between those mean-subtracted traces. Note that the same analysis based on responses to drifting gratings data gave consistent results (**Figure S4**).

### Statistical analyses

All data are expressed as the mean ± standard error of the mean (SEM), unless stated otherwise. The two-sided Wilcoxon signed-rank test was used to compare two paired groups. Friedman test was performed when more than two groups were compared. When significant, it was followed by Tukey-Kramer method. For distribution of preferred orientation and direction, Kolmogorov-Smirnov test was used. For data from slice whole-cell patch clamp recordings, Wilcoxon rank-sum test was used to compare two groups. Fisher’s exact test was used to compare the proportions of connected cell pairs. Kruskal-Wallis test was used when more than two groups were compared. The variances between groups were assumed to be similar. Throughout the study, *P* < 0.05 was considered statistically significant, other than the definitions of visually responsive and selective cells (see Data analysis of visual response). No statistical methods were used to pre-determine sample sizes, but our sample sizes are similar to those generally employed in the field.

## DATA AND SOFTWARE AVAILABILITY

All data and analysis code are available upon reasonable request to the Lead Contact, Kenta M. Hagihara.

---

**Figure S1.**
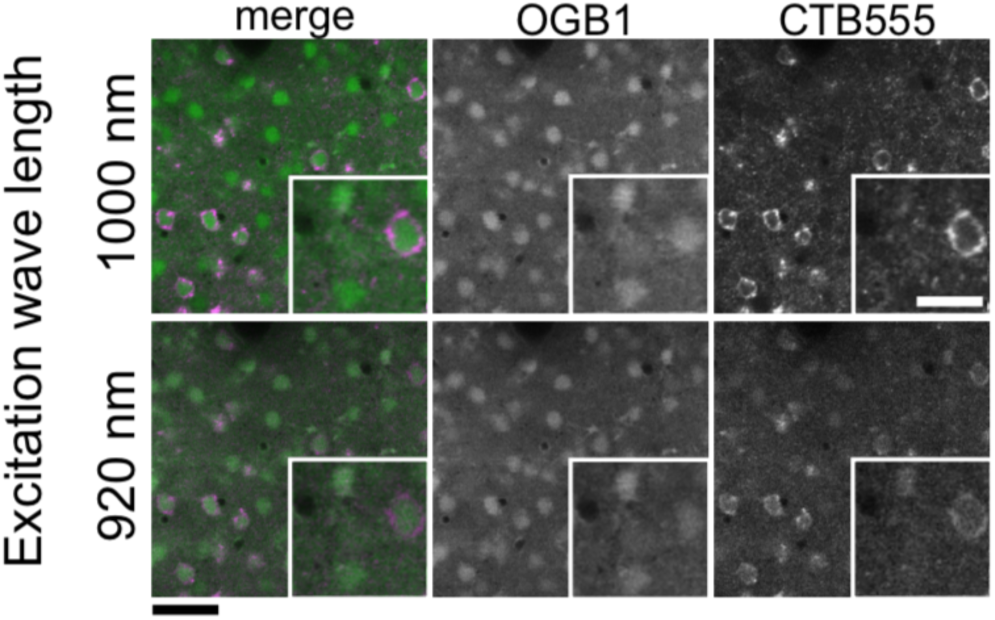
Identification of CPNs *in vivo*, Related to Figure 1. Two-photon imaging of CPNs with a 1000 nm or 920 nm excitation laser. Green: OGB1; Magenta: CTB555; Scale bar: 50 μm (main panel); 10 μm (inset)

We realized that the OGB1 signal had contaminated the CTB555 signal when we imaged with a 920 nm excitation laser, which (or similar wavelength lasers) has conventionally been used in previous studies to image calcium indicator-labeled neurons (Chen et al., 2013; Li et al., 2015). However, with a 1000 nm laser, the signal contamination was largely reduced, and we could reliably identify CPNs by visual inspection. Thus, we used a 1000 nm laser for excitation of both OGB1 and CTB555 in this study.

**Figure S2.**
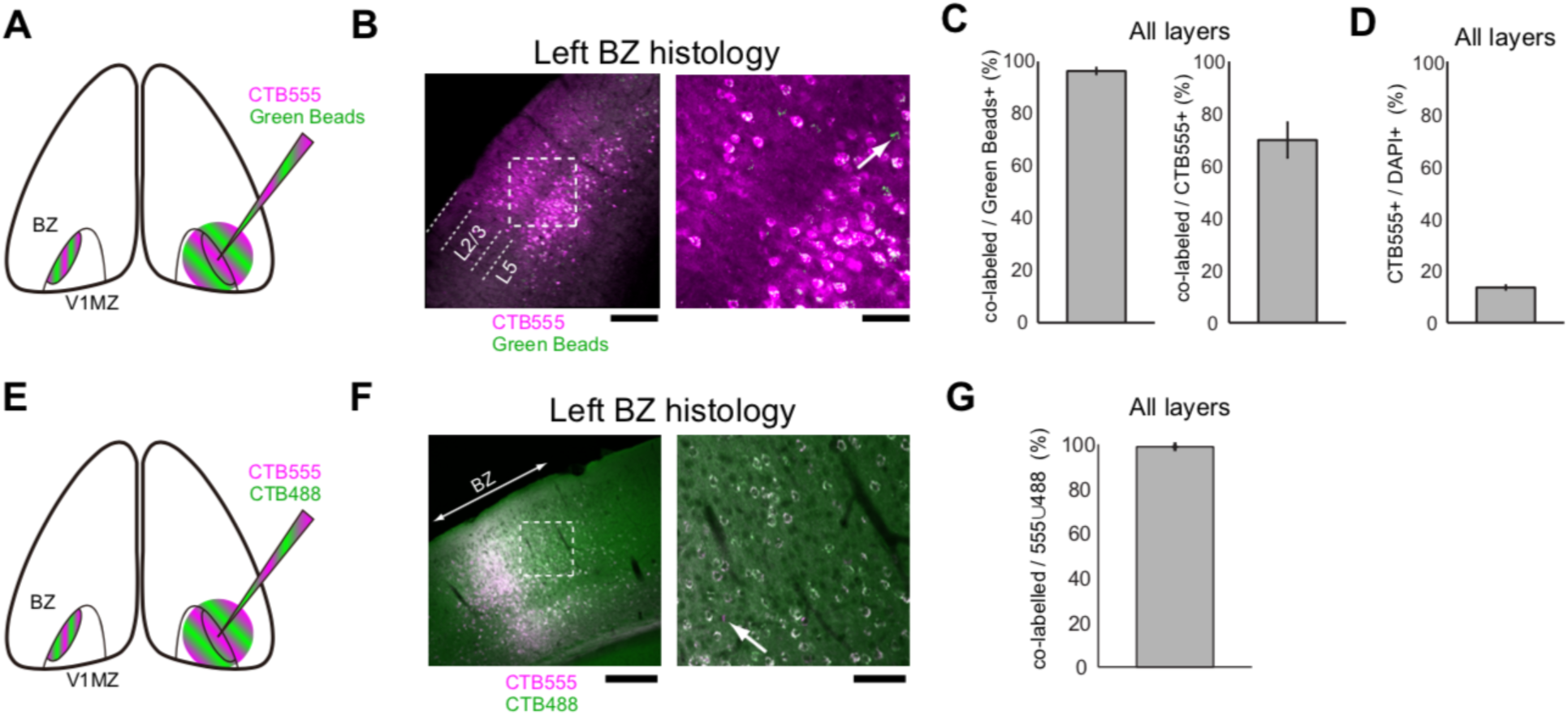
Assessment of the efficiency of retrograde-labeling with CTB, Related to Figure 1. **A.** Schematic drawing of co-injection of CTB555 and Green Beads. **B.** A histological coronal section of the BZ in the contralateral hemisphere to that of the injection site. (Left) The area surrounded by the dashed lined box indicates the area shown in the right panel. Scale bar: 200 μm; (Right) The arrow indicates a neuron labeled only with Green Beads. Scale bar: 50 μm; **C.** Quantification of the co-labeled neurons. **D.** Quantification of the CTB555-positive neurons as a percentage of DAPI-positive cells. **E.** Schematic drawing of co-injection of CTB488 and CTB555. **F.** A histological coronal section of the binocular zone (BZ) in the opposite hemisphere to the injection site. (Left) The area surrounded by the dashed box indicates the area shown in the right panel. Scale bar: 250 μm; (Right) The arrow indicates a neuron labeled only by CTB555 (magenta). Scale bar: 50 μm; **G.** Quantification of the co-labeled neurons. The efficiency of CTB488 and CTB555 can be regarded as almost 100% in our system. This result confirms that neurons not labeled with CTB555 can be largely regarded as non-CPNs in the current study.

To characterize the efficiency of our CPN labeling, we injected CTB555 together with another retrograde tracer Green Beads, into the BZ, and calculated the percentage of neurons that were labeled with both CTB555 and Green Beads when compared with all the Green Beads positive neurons. We found that almost all Green Beads-labeled neurons were also positive for CTB555 (the percentage of double-positive cells was 96.3 ± 1.2% (mean ± SD), n = 3 mice), while a very small number of neurons showed only Green Beads fluorescence (**Figure S2A–C)**. This suggests that CTB555 is taken up so efficiently that there would be few false negatives, if any, in our CPN labeling (see **Methods** for the rationale), as previously described in a different projection circuit (Bortone et al., 2014). In addition, by calculating the fraction of CTB555-positive neurons among DAPI-positive neurons, 13.5 ± 0.8% (mean ± SD) neurons were identified as CPNs (**Figure S2D**). Consistently, dual injection of CTB555 and CTB488 revealed that their efficacy is quite high in our system (**Figure S2E–G**).

**Figure S3.**
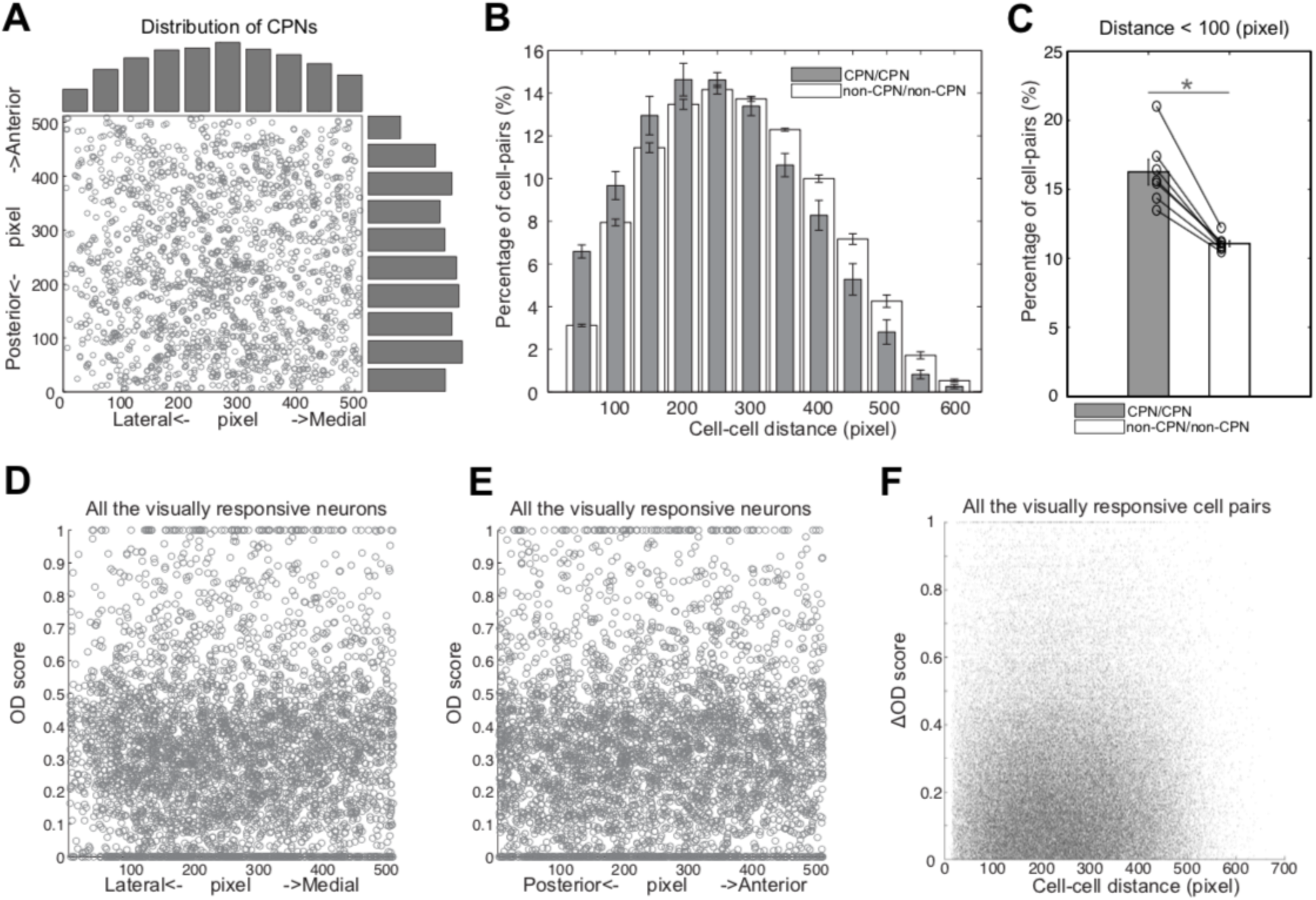
Spatial organization of CPNs and spatial representation of eye preference, Related to Figure 2. **A.** Scatter plots for locations of CPNs. The outside histograms show their distribution along the lateral-medial axis and the anterior-posterior axis. **B.** Distribution of cell-cell distance between CPNs and between all neurons. **C.** Proportion of cell pairs with a distance below 100 pixels. *: *P* < 0.05; (Wilcoxon signed-rank test) **D, E**. Scatter plots for ODscore against the location of the cell in the lateral-medial axis (**D**) and in the anterior-posterior axis (**E**). **F**. Scatter plot of the difference of OD scores (ΔODscore) against the cell-cell distance of all the visually responsive cell pairs.

To study whether CPNs show specific spatial organization, we plotted locations of CPNs in the visual cortex. We found that CPNs were more frequently located at the center of imaging ROIs along the medial-lateral axis and showed gradual tails at the periphery (**Figure S3A**). Together with the fact that OGB1 injection was performed to largely cover the CTB555 positive region, CPNs most likely show a gradual distribution, with a higher density at the border between V1 and the lateral secondary visual cortex, and this gradual distribution contributed to the slightly left-shift in cell-cell distance distribution of CPNs compared to that of non-CPNs (**Figure S3B, C**, *P* = 0.016, *n* = 7 mice). Thus, it is unlikely that mouse visual cortex has microstructures of CPNs in the BZ as found in the Long Evans rat (Laing et al., 2015) or more higher mammals, rather it shows gradual spatial organization.

When we plotted the ODscore against the position in the lateral-medial and anterior-posterior axes of the visually responsive neuron, we did not see any evidence of a microstructure (**Figure S3D, E**). In addition, plotting the difference of OD scores (ΔODscore) against cell-cell distance did not show any clustering (**Figure S3F**). As such, and consistent with previous reports on rodents (Antonini et al., 1999; Drager, 1974; Gordon and Stryker, 1996; Mrsic-Flogel et al., 2007; Scholl et al., 2015) (but see (Laing et al., 2015) for results in the Long Evans rat), we found little evidence for a clear columnar organization of neurons with similar ODscores. Instead, the BZ in the mouse is likely to be uniformly organized in terms of ocular dominance response properties.

**Figure S4.**
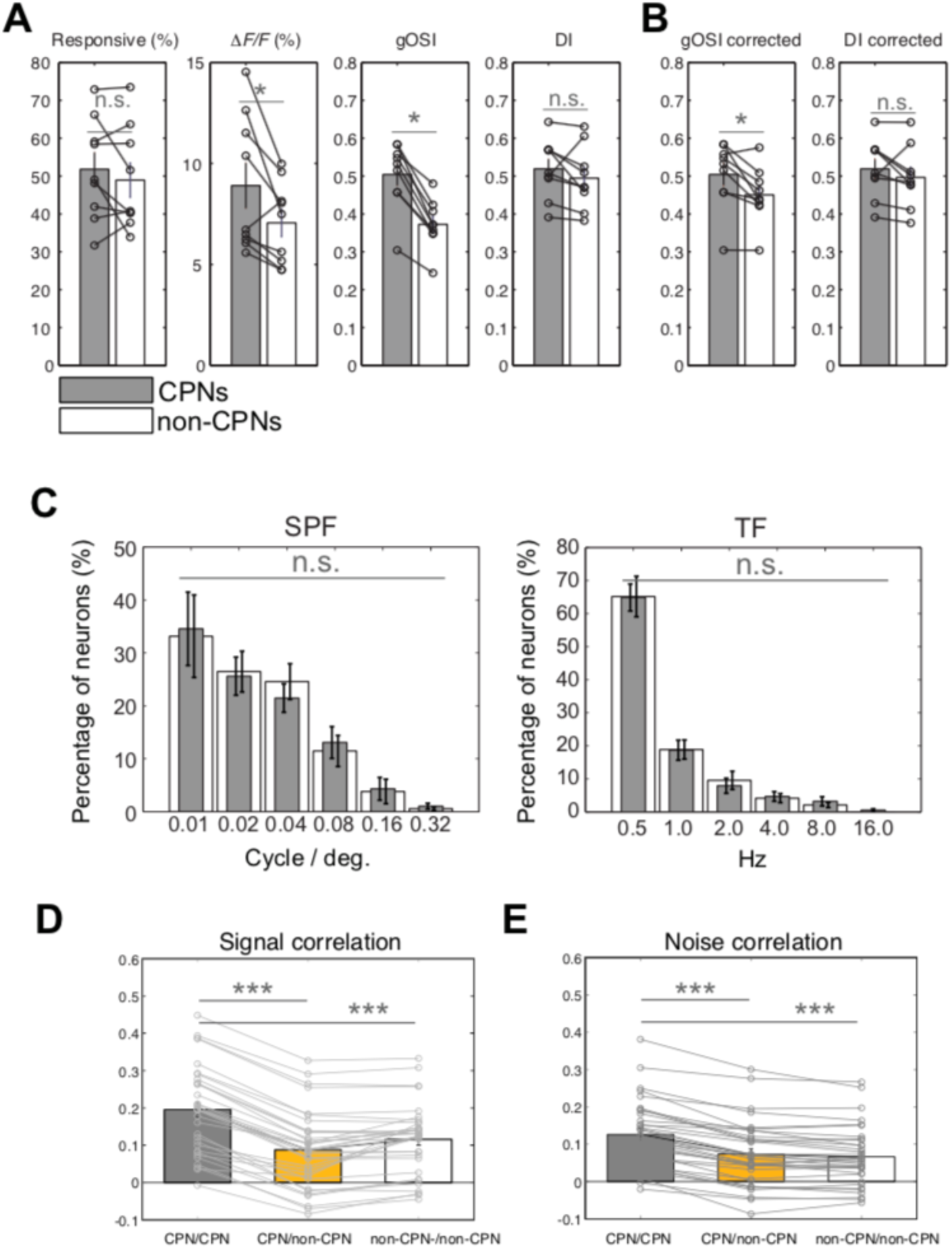
Response properties of CPNs and non-CPNs to drifting gratings, Related to Figure 2. **A.** Responsiveness, ΔF/F, gOSI, and direction index (DI) of CPNs and non-CPNs. *: *P* < 0.05; n.s., non-significant; (Wilcoxon signed-rank test). **B.** gOSI and DI of CPNs and inhibitory neuron-corrected non-CPNs. **C.** Distribution of preferred SPF and preferred TF of CPNs and non-CPNs. n.s., non-significant; (Friedman test). **D, E**. Means of signal (**D**) and noise (**E**) correlation values from each plane based on 12-direction drifting grating stimuli. ***: *P* < 1.0 × 10^−3^; (Friedman test followed by Tukey-Kramer).

Consistent with the results using ocular dominance stimuli, there was no significant difference in the fraction of responsive neurons between CPNs and non-CPNs (**Figure S4A**, 51.8 ± 4.5%, 48.9 ± 4.6%, *P* = 0.65; Wilcoxon signed-rank test, *n* = 9 mice). In addition, we found a slightly larger ΔF/F amplitude in CPNs than in non-CPNs (**Figure S4A**, *P* = 0.039). We found no significant difference in the distribution of preferred orientation and direction between CPNs and non-CPNs (*P* > 0.5, Kolmogorov–Smirnov test, *n* = 9 mice), indicating that information concerning each orientation and direction is evenly transmitted to the other hemisphere. A higher tuning sharpness was indicated for CPNs, which showed a larger gOSI (see **Methods**), reflecting orientation tuning sharpness, than for non-CPNs (*P* = 0.004; Wilcoxon signed-rank test) (**Figure S4A**), while there was no difference in DI, which reflected direction tuning sharpness (*P* > 0.05). We also studied spatial and temporal frequency (SPF/TF) tuning of CPNs and non-CPNs and found no difference in the distribution of preferred SPF and TF between CPNs and non-CPNs (**Figure S4C**, *P* > 0.8, *n* = 10 mice; *P* > 0.8, *n* = 8 mice; Friedman test).

The difference in gOSI could be attributed to the lower gOSI of inhibitory neurons (Hofer et al., 2011; Kerlin et al., 2010; Sohya et al., 2007), because non-CPNs comprise both excitatory and inhibitory neurons, while CPNs comprise only excitatory neurons. To exclude this possibility, neurons with the lowest 25% gOSI were considered to be putative inhibitory neurons and were eliminated from non-CPNs (Hagihara et al., 2015). The remaining populations were regarded as excitatory neurons, and compared with CPNs. Given that the percentage of inhibitory interneurons is approximately 20%, and that a small fraction of inhibitory interneurons shows a high gOSI (Hofer et al., 2011; Kerlin et al., 2010; Sohya et al., 2007), this correction could be slightly too strict. However, even with this potential over-correction, CPNs continued to show a slightly but significantly higher gOSI than non-CPNs (**Figure S4B**, *P* = 0.019). Overall, CPNs are likely to be more sharply tuned than non-callosal projection excitatory neurons.

**Figure S5.**
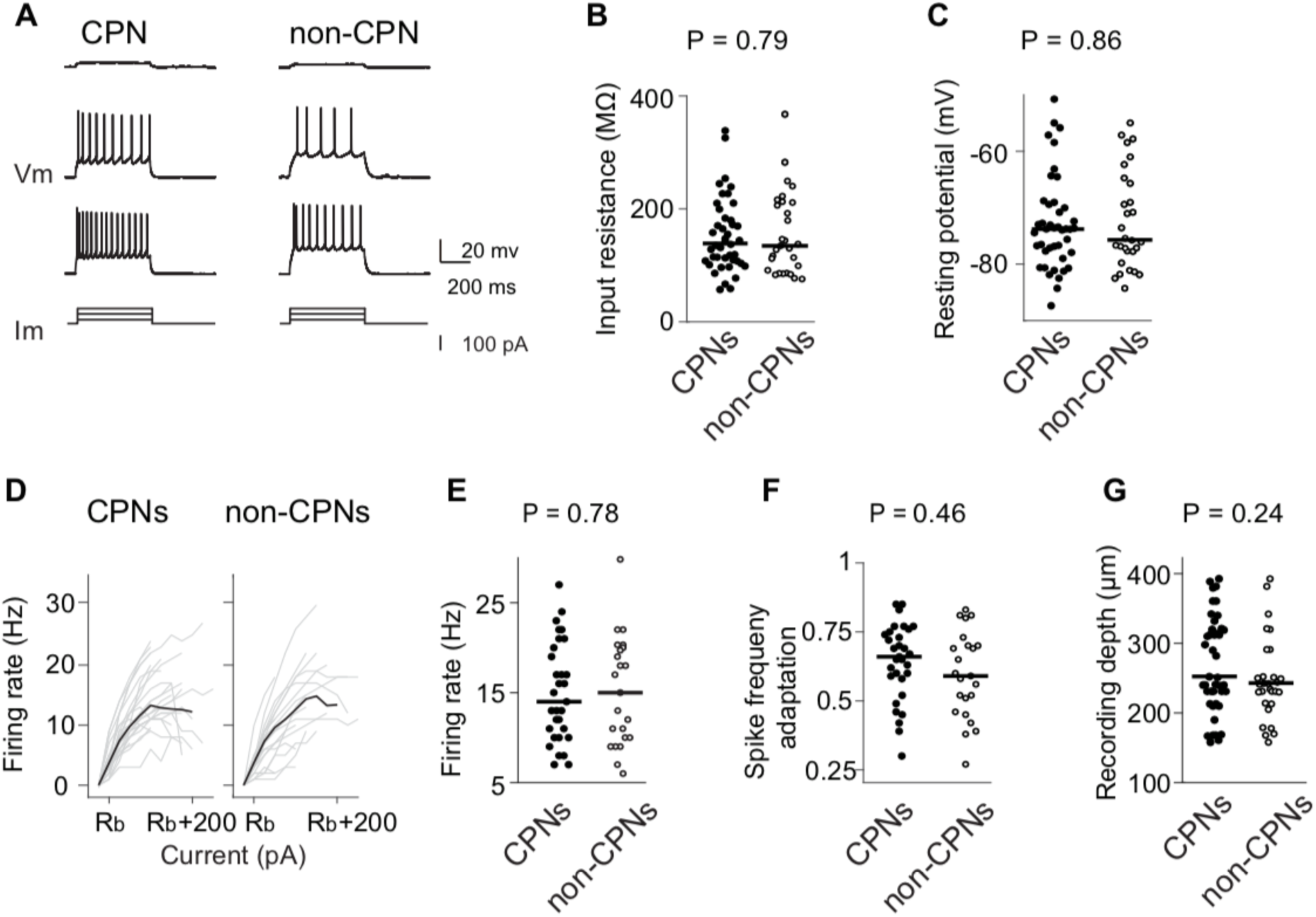
Intrinsic membrane properties of CPNs and non-CPNs, Related to Figure 3. **A.** Example membrane voltage responses to current injections for CPN (left) and non-CPN (right). Step current injections shown in lower panel and corresponding voltage responses in upper panel. **B.** Input resistance determined by the slope of the regression line between the hyperpolarizing current pulse (from −50 pA to 0 pA in 25 pA steps) and peak voltage responses during the first 50 ms after current onset. Circles indicate values obtained from individual neurons and the lines indicate the median (CPNs, n = 43; non-CPNs, n = 29). **C.** Resting membrane potential determined just after establishment of whole-cell configuration (CPNs, n = 43; non-CPNs, n = 29). **D.** The current intensity - firing frequency relationship (I-F relationship) above the rheobase (Rb) current for CPNs (left) and non-CPNs (right). Each thin line indicates individual I-F curves. Thick lines indicate the averaged I-F curves. The Rb was defined as the lowest current intensity (25 pA step with 1 s duration) to induce at least one action potential. **E.** Maximum firing frequency for CPNs (n = 31) and non-CPNs (n = 23). **F.** Spike frequency adaptation defined as the ratio of the first and the last inter-spike intervals of the spike train evoked at current adjusted for each neuron to induce its maximum firing frequency (CPNs, n = 31; non-CPNs, n = 23). **G.** Recording depth from pia surface for CPNs and non-CPNs (CPNs, n = 43; non-CPNs, n = 29). The *P* value was calculated by Wilcoxon rank-sum test.

